# Inferring the Genetic Influences on Psychological Traits Using MRI Connectivity Predictive Models: Demonstration with Cognition

**DOI:** 10.1101/777821

**Authors:** Alexander S. Hatoum, Andrew E. Reineberg, Philip A. Kragel, Tor D. Wager, Naomi P. Friedman

## Abstract

Genetic correlations between brain and behavioral phenotypes in analyses from major genetic consortia have been weak and mostly non-significant. fMRI models of systems-level brain patterns may help improve our ability to link genes, brains, and behavior by identifying reliable and reproducible endophenotypes. Work using connectivity-based predictive modeling (CBPM) has generated brain-based proxies of behavioral and neuropsychological variables. If such models capture activity in inherited brain systems, they may offer a more powerful link between genes and behavior. As a proof of concept, we develop models predicting intelligence (IQ) based on fMRI connectivity and test their effectiveness as endophenotypes. We link brain and IQ in a model development dataset of N=3,000 individuals; and test the genetic correlations between brain models and measured IQ in a genetic validation sample of N=13,092 individuals from the UKBiobank. We compare an additive connectivity-based model to multivariate LASSO and ridge models phenotypically and genetically. We also compare these approaches to single “candidate” brain areas. We find that predictive brain models were significantly phenotypically correlated with IQ and showed much stronger correlations than individual edges. Further, brain models were more heritable than single brain regions (h^2^=.155-.181) and capture about half of the genetic variance in IQ (rG=.422-.576), while rGs with single brain measures were smaller and non-significant. For the different approaches, LASSO and Ridge were similarly predictive, with slightly weaker performance of the additive model. LASSO model weights were highly theoretically interpretable and replicated known brain IQ associations. Finally, functional connectivity models trained in midlife showed genetic correlations with early life correlates of IQ, suggesting some stability in the prediction of fMRI models. We conclude that multi-system predictive models hold promise as imaging endophenotypes that offer complex and theoretically relevant conclusions for future imaging genetics research.

---

The vulnerability and advantages conveyed by genetics via behavior is likely due to individuals inheriting particular brain systems that support behavior. Perhaps this is why genetics and MRI are the two most utilized biostatistical approaches for studying psychological phenomena. Luckily, these two fields are developing in parallel, as both whole-genome analyses^1, 2^ and multivariate brain analyses(i.e. including multiple brain areas or connections simultaneously)^3^ are expanding the inferences and predictions we can make about psychological traits. As both fields expand, our ability to translate across biological psychology will depend on our ability to integrate the explosion of techniques in both fields.

In genetics, the standard discovery procedure is a genome-wide association study (GWAS), which requires large (10’s-100’s of thousands) samples and is agnostic to *a priori* hypothesized SNP associations. Further, these whole-genome techniques can be used to give us a measure of the genetic correlation between phenotypes, or the degree to which two traits share genes and are likely inherited together.

However, the effects of individual genetic variants in GWAS are incredibly small. Given these small effect sizes and the high threshold necessary for multiple testing correction across the genome (alpha = 5×10^-8^), GWAS studies have necessarily adopted a coarse phenotype approach to enable larger sample sizes. We are now beginning to understand the consequences of this necessary evil, as coarse phenotypes can make results difficult to interpret and can lead to less specificity in understanding behavioral traits, as some have demonstrated with depression^4^. Further, poor phenotypes limit the types of conclusions we can make from genetic correlations and whole-genome patterns.

One early suggested approach to improve genome-wide discovery and theoretical interpretation is to use intermediary traits, or endophenotypes: traits that are more closely related to the genetic expression of a distal behavioral outcome and likely to be more heritable^5^. For example, hippocampal volume is decreased in individuals with schizophrenia and non-affected related family members^6^, and likely closer to genetic inheritance than schizophrenia. Thus, it was thought that a GWAS of hippocampal volume should capture (some of) the genes influencing schizophrenia.

For years, the endophenotype approach presented an attractive alternative to the coarse approach. The most ubiquitous endophenotype for psychological and neurological phenomena is the brain or imaging genetics studies. Figure 1 shows a chart of the terms “endophenotype” and “imaging genetics” that have appeared since 2010, showing thousands of articles published on the topic each year. Further, imaging is a useful foray in the endophenotype literature as large GWAS cohorts are being assembled for MRI imaging phenotypes and offers some insight into the relative success of endophenotypes to date.

**Figure 1.**
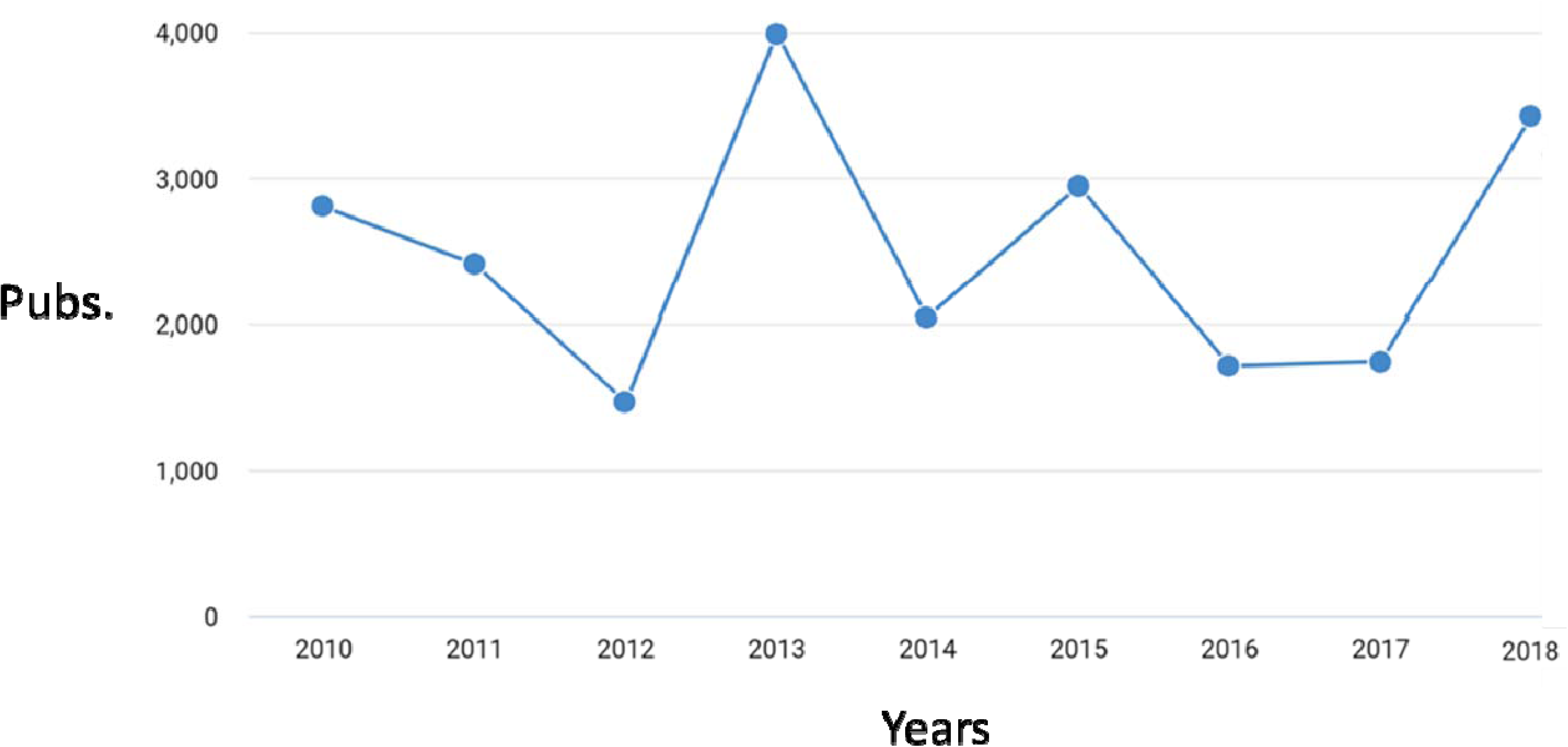
Rate of words “Imaging Genetics” and “Endophenotype” appearing together by Google Scholar search across 8 years. Pubs.=Publications from Google scholar search.

The two goals of endophenotypes were to help discover new variants and explain the mechanisms of genetic inheritance. Unfortunately, these two goals of endophenotypes in imaging genetics have been unrealized. No new genetic variants have been discovered due to imaging endophenotypes (yet)^7^. Further, genetic associations between well-known brain endophenotypes and traits are low and non-significant in the largest and most powerful studies to date^8^, weakening the claim that current endophenotypes are explanatory tools for scientific theory.

We argue that the success of the imaging endophenotype approach can be improved by considering a more integrative and predictive multi-system brain approach to endophenotypes. We hypothesize that potentially including areas from across the whole-brain will capture (more of) the genetic variance underlying behavioral phenotypes than a single brain measure. For example, although the hippocampus has been consistently associated with schizophrenia, there was little success in finding genetic associations between hippocampal volume and schizophrenia^8^. If genes leading to the development of schizophrenia represent the inheritance of brain systems, then integrating information across the brain could increase our ability to find these genetic associations, i.e. not just including the hippocampus, but the hippocampus likely works in a system with other brain areas that more reliably predicts schizophrenia, and it is that multi-system level prediction that should be utilized in imaging genetics studies.

## Multi-system brain predictive models vs. candidate approaches

While there are numerous neuroimaging brain-to-outcome associations in the literature, many of these have failed to make an impact on clinical practice. Woo, Chang, Lindquist, & Wager (2017)^9^ offered a possible solution through the use of multi-system brain predictive modeling. These approaches are preferred to those focused on single candidate brain regions because they are more powerful omnibus summaries for the whole-brain, allow for flexible tests of reproducibility, incorporate more information, and are more predictive than standard approaches^10^.

Thus, with the current approach in the literature, specific associations are found and then established as endophenotypes based on univariate tests of their association with the outcome and in related family members, akin to a “candidate endophenotype” method. It is this approach that has failed to live up to the expectations of the endophenotype claims, and is not fully utilizing new trends in the imaging literature. We argue that the prediction by these multi-system brain models can be utilized as a GWAS phenotype. In other words, we propose “multi-system brain endophenotypes” as new path forward in the gene discovery literature. Specifically, we call the prediction by multivariate approaches that consider many brain regions simultaneously “Multi-system brain endophenotypes”.

While complete literature reviews of these multi-system brain models are available elsewhere^3, 11^, models of outcomes such as pain responsivity^12^ sustained attention^13^ and autism^10^, have been developed and replicated across large cohorts. Any of these models and numerous others could offer a theoretically relevant endophenotype useful for GWAS studies. Finally, while these phenotypes have statistical and theoretical advantages to standard approaches, they give the added advantage of offering many new phenotypes for GWAS discovery than those measured; for example, a GWAS of sustained attention patterns may inform understanding of ADHD.

## Connectivity-Based Predictive Modeling (CBPM)

Brain imaging has many measurement approaches, or “modalities”, that can be used to generate measures of the brain, e.g., aspects of anatomical size or flow of oxygenated blood in the brain. Thus, a first step in developing a multi-system brain model is the choice of brain data. For our demonstration, we used functional connectivity as measured by resting-state fMRI, a method that quantifies variation in the degree to which brain regions intrinsically correlate in the time course of the blood-oxygen level dependent (BOLD) signal while individuals are not overtly directed toward a particular task set (i.e., are at rest)^14^. A key element of how CBPM is implemented using functional connectivity is the scale at which connectivity is considered – typically, all pairwise connections between 10s to 100s of regions are considered.

## This Study

To demonstrate the effectiveness of multi-system brain predictive models as phenotypes for genetic research, we generate CPBMs of intelligence (IQ) and estimate the genetic correlation between brain-based predicted IQ and measured IQ. Specifically, for predictive models to be useful endophenotypes, they must be associated with the trait of interest^5^ and be theoretically relevant, interpretable, and generalizable^15^ at the genetic level. Figure 2 shows our workflow through several analyses that help to achieve this aim, along with the research question answered by each analysis.

**Figure 2.**
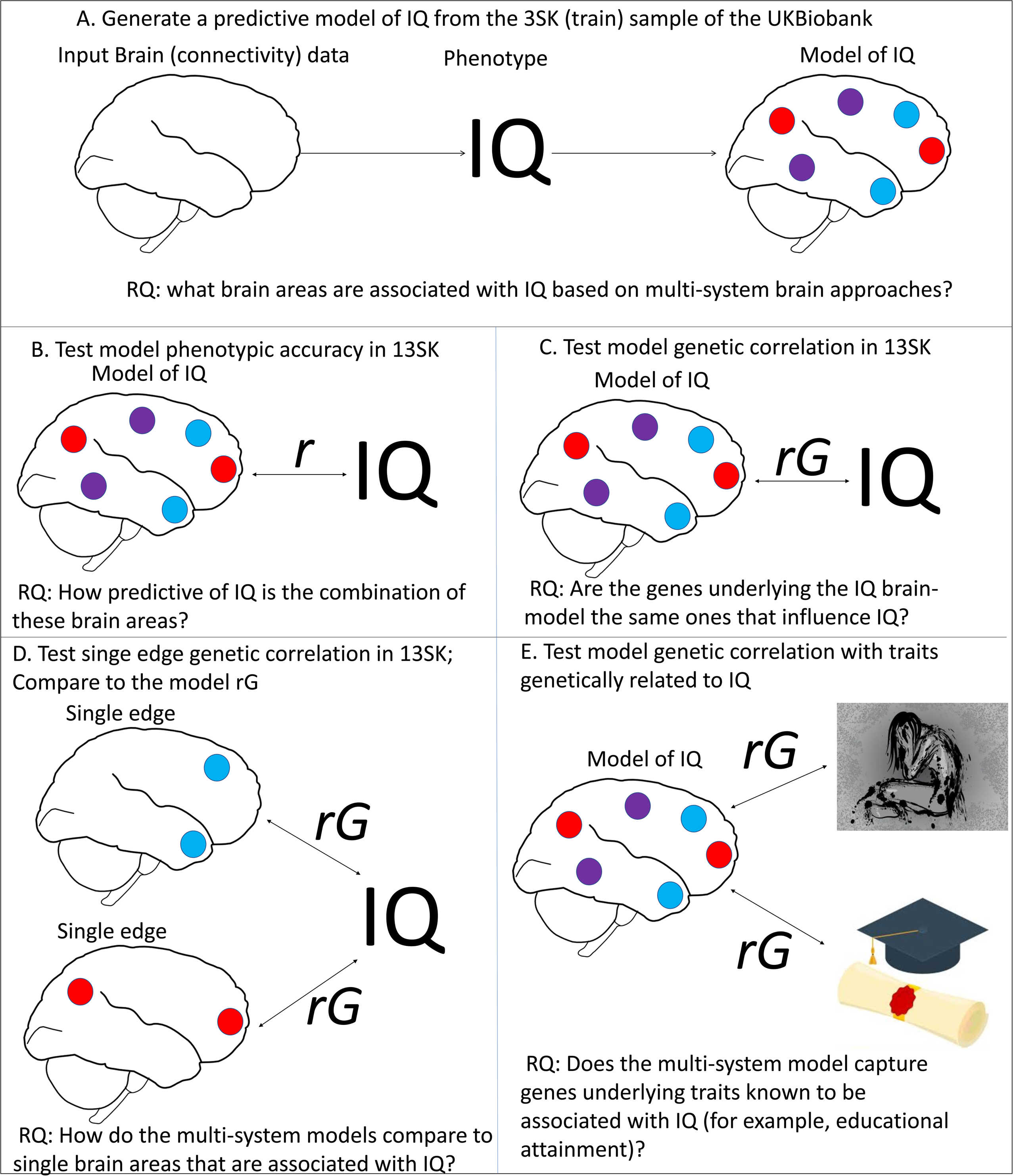
Workflow for main analyses in this study. First (A) we generated connectivity-based predictive models (CBPM) of IQ in a sample of 3,000 individuals with brain data from the UKbiobank. Next (B) we confirmed the out-of-sample predictive accuracy of these models phenotypically by relating brain-based predicted IQ from brain data with measured IQ in a genetic validation set of ∼13,000 individuals of European descent. Because the sample of 13,000 individuals also had genetic data, (C) we tested the genetic correlation between predicted IQ and measured IQ in this sample. Next, (D) we ran the genetic correlation between the strongest single edges and IQ and compared them (using the 95% confidence intervals) to genetic correlations from multi-system brain brain-based predicted IQ. Finally, (E) to confirm that these genetic correlations were capturing relevant theoritical aspects of IQ, we estimated the genetic correlation between brain-based predicted IQ and known genetic correlates of IQ.

First, (A) in a sample of 3,000 individuals from the UK biobank we develop three predictive models of IQ with three different predictive techniques. We extract the weights from the models to see how they align with past connectivity-based studies of IQ. Next, (B-C) to test the effectiveness of the prediction of these three models as endophenotypes, in a genetic validation sample of 13,092 unrelated individuals we estimate the phenotypic and genetic correlation between predicted and measured IQ and conduct a GWAS of each model’s prediction and interpretability. In this case, we use the larger sample as validation to have more power to estimate genetic correlations and more power for comparison of multi-system brain measures and single brain measures. After estimating phenotypic and genetic correlations between brain-based predicted IQ and measured IQ, we (D) compare the multi-system brain models predictions to the phenotypic and genetic associations between IQ and single associated edges. Finally, (E) we relate our predicted model of IQ to behaviors known to be genetically correlated with IQ.

We should note that the purpose of this work is not to test all possible machine learning methods and relevant applications (we only look at three here), but rather to create a very simple procedure for others to follow. We also report two post-hoc analyses that should help develop this area of research for future genetic studies. Specifically, we use learning curves to demonstrate the improved accuracy of multi-system predictive brain models and demonstrate what sample sizes will likely be needed for this approach in the future. We then use lesion analysis to expand the theoretically relevant neurological implications of edges discovered by our machine learning approaches.

## Method

### Participants

The UKBiobank is a large population cohort of over 500,000 individuals, aged 36-72. We restricted our analysis to 16,092 individuals who had MRI data, IQ scores, and who were of European descent. We split the sample (randomly) into a sample of 3,000 (S3K) individuals (mean age = 55.07, standard deviation age =7.42) and 13,092 (S13K) individuals (mean age=55.12, standard deviation age = 7.45), for different aspects of the project. Though non-intuitive, we chose to use the S13K individuals as a genetic validation set as this sample size had appropriate power to detect genetic correlations^16^. Further, the sample of 3,000 is orders of magnitude larger than most samples previously used in MRI research to develop predictive models^11, 13^. Because inference about the performance of these techniques would benefit from larger sample sizes, to estimate learning curves at increasing sample sizes (phenotypically), we use S13K to estimate the model and S3K to test the three learning techniques phenotypically in a post-hoc analysis to help others establish appropriate sample sizes for training and test sets in future phenotypic work.

IQ in the UKBiobank was measured at four time points: three in-person assessments and an online assessment. All assessments of IQ were two-minute tests of fluid intelligence: Participants had 2 minutes to answer as many questions as possible in a sequence of 13 questions. We estimated a latent factor model across all four time points in the full sample of ∼500,000 individuals and extracted the factor score as a measure of IQ to increase reliability due to the short assessments. Data from individuals who “abandoned” the later time points were treated as missing in factor analysis. Factor scores from the imaging sample are used in this analysis.

### Brain data: Parcellation and Measurement

The UK Biobank project provides highly summarized rs-fMRI data in the form of 2 full connectivity matrices per participant: pairwise correlations between 25 and 100 regions, as derived from large independent components analyses (ICAs) conducted with FSL’s MIGP-Melodic tool. These correlations within each individual represent the training features/variables of analysis, typically called “edges.” Because these 25- and 100-dimension parcellations include noise components, the actual number of signal components is lower than 25 and 100. We chose the 100-dimension parcellation, which was reduced to 55 signal components based on an analysis by the UKBiobank imaging group. The resulting data, per participant, was a full 55 x 55 correlation matrix, leaving us with 1485 unique functional connections/edges after excluding the diagonal. Each edge was residualized on age and sex before training.

### SNP data processing and associations

We used UKBiobank participants whose data were imputed to the Haplotype Reference Consortium (see McCarthy et al. 2016 Nature Genetics for more)^17^, 1000 Genomes, and UK10K reference panels by the UK Biobank^18^ and of European descent. Subjects were genotyped on a UK BiLEVE array or the UKBiobank axiom array. After removing individuals with mismatched self-reported and genetic sex, we filtered imputed SNPs using a Hardy-Weinberg equilibrium P value threshold of >1×10^6^, variant missingness > 0.05, imputation quality score (INFO) >0.95, and minor allele frequency (MAF) above 0.01, retaining 7,391,068 SNPs. More information is available in a prior publication^18^.

### Training procedures

We generated three scores for multi-system brain predictive models. First, we adapted the Shen et al. 2017^14^ CBPM procedure, which considers each edge of the connectome as an individual additive predictor. The procedure sums positively and negatively associated edges (below an associated p-value threshold) and uses the negative and linear sum scores as separate beta weights in a linear model to predict the outcome of interest. We test the Shen et al. model at p-value cutoffs of .05 in initial analysis, as this is the typical value chosen in the literature^13^. In the post-hoc analysis, we use learning curves to demonstrate how adjusting the p-value threshold changes the prediction, if at all.

The second and third models were based off the procedure from Tobyne et al. 2018^19^ and Michel et al. 2012^20^, in which cross-validated regularized regression was used to train CBPM models. We trained ridge regression (a.k.a L2 regularization) and LASSO regression (Least Absolute Shrinkage and Selection Operator, a.k.a L1 regulations) with 10-fold validation (used to choose the parameter values and regularization parameters), implemented in the R package glmnet^21^. We conducted a grid search of values from 0 to 10,000 (default of our package) for optimal regularization parameters (called “Rho” or “Lambda”) for each model. To interpret the predictive models, we extracted the weights for each edge and from each model. We then plotted the weights in matrix form with a set of superordinate resting-state network assignments (from Yeo et al.)^22^ annotated along the axes to increase interpretability of the results, given high familiarity with this very popular network parcellation.

To evaluate whether certain functional connections were necessary for successful prediction, we additionally compared the LASSO and ridge regression models. This comparison evaluated whether IQ is better predicted by differences in the connectivity of a few brain regions, as characterized by LASSO regression, or is better reflected by smaller changes in the connectivity between multiple brain regions, as modeled by ridge regression. Specifically, to evaluate whether the regions identified by the LASSO model were driving the multi-system brain signal in the ridge model, the key connections identified by fitting the LASSO model were “lesioned” from the full connectivity matrix, and the remaining connections were used to fit the ridge regression model. We conducted a corrected repeated *k*-fold cross-validation test (10 iterations of 10-fold cross-validation; see ^23^), to estimate whether LASSO or ridge regression performed better.

To see how effective these models are at predicting IQ, in the training 3SK sample we trained a ridge regression, LASSO Regression, and Shen CBPM and then used the weights extracted from the 3SK sample to predict IQ in 13SK genetic validation sample. After estimating the scores, we used a Pearson’s correlation (instead of mean-squared error) to determine phenotypic association between brain-based predicted IQ (from the models’ summary score) and measured IQ, as we thought Pearson’s r would be most comparable to the genetic correlation (rG).

### Test of scores as GWAS proxies for measured phenotypes

#### Univariate heritabilities

In the 13SK sample, we used a mixed-effects model procedure through BOLT-LMM^24^ to estimate the univariate heritability’s of brain-based predicted IQ scores for the ridge regression, LASSO regression, and Shen et al. procedure in the out of sample predictions. We also compared them to the heritability of measured IQ in the 13SK sample.

#### Bivariate heritabilities

To determine if the GWAS proxies were capturing the same genes as measured IQ, we estimated the genetic correlation between measured IQ and out-of sample brain-based predicted IQ in the 13SK genetic validation data set. To do so, we used Bolt-LMM to estimate the genetic correlations between all three brain-based predicted IQ scores and measured IQ, separately.

To compare the genetic correlations between multi-system brain models and individual brain connections, we ran a test of association (controlling for sex and age) in the 3SK sample of each edge and IQ. After Bonferroni correction, 6 individual edges were significantly associated with IQ (Table 2) in the 3SK. We then estimated the genetic correlation between each of these 6 edges and IQ in the 13SK. We compare these to the multi-system brain correlations and correct for multiple testing in the genetic correlations (9 tests) using a Bonferroni correction.

#### GWAS

To see if these scores yield useful GWAS discoveries we then used BOLT-LMM to conduct a mixed-effects GWAS (with leave one chromosome out to keep from overfitting our GRM per SNP) for IQ and each predictive model in the 13SK sample. We chose BOLT-LMM to increase power and account for the polygenic background of our trait^24^. For each GWAS, we examined a Manhattan plot of individual SNP p-values and also explored the observed p-values deviation from expected p-values using QQ plots.

#### Genetic correlations with IQ covariates

Finally, to see if brain-based predicted IQ from functional connectivity showed similar patterns of genetic correlation as IQ, we tested genetic correlations between summary statistics of the GWAS for each IQ prediction (Ridge, Lasso, and Shen CBPM) and related traits through the LDhub^16, 25^. We chose 8 traits that have been previously shown to be genetically correlated with IQ in a major meta-analysis^26^ and were theoretically relevant to IQ. We considered IQ proxies (educational attainment^27^, and childhood IQ^28^), or psychiatric covariates of IQ (autism and depressive symptoms^27^), anthropometric traits (infant head circumference^29^, height^30^), and evolutionary linked traits (age of first birth^31^), as these are the typical types of hypotheses tested using genetic correlations between IQ and a covariate of IQ. We did not consider the schizophrenia and IQ genetic correlation because the schizophrenia sample that was publicly available at time of analysis includes mixed ethnicity. We also did not run exhaustive LD-score correlations between brain-based predicted IQ and all trait summary statistics available online to reduce the number of tests.

### Testing generalizability of brain predictive models for IQ

To test effective sample sizes for out of sample prediction of IQ, we estimated learning curves for each algorithm as a function of sample size. To preface results, we chose to do this because more predictive models were also more genetically associated with measured IQ. We plotted the canonical correlation between brain-based predicted IQ and measured IQ in both the training set and test set, increasing sample sizes of 50 to 13092 individuals in increments of 50. We used the S13K as the training and S3K as the validation set for this analysis to increase the spectrum of sample sizes tested. We estimate curves for the (1) ridge, (2) LASSO, (3) Shen et al. at p-value threshold of .005, (4) p-value threshold of .01, (5) p-value threshold of .05, (6) p-value threshold of .10, and (7) no p-value threshold (summing up all edges). Importantly, this is the first work to compare the Shen et al. predictive models at varying p-value thresholds.

## Results

### Phenotypic prediction

We began by training our models in the S3K sample and testing their predictive accuracy in the S3K and out-of-sample S13K. Table 1 shows the results from the phenotypic prediction in both samples; with effectiveness represented by the correlation between measured and brain-based predicted IQ. All three models showed significant phenotypic prediction in the training and test set. Phenotypic prediction in the training set varies between r = .322 - .476. In the test set, the prediction ranged between r = .187 - 212. In both the train and test set the ridge regression was the most predictive model (post-hoc analysis will show how model prediction was affected by our sample size). The test set may be lower because cross-validation is not a panacea for overfitting as cross-validation still capitalizes on stochastic error^32^.

**Table 1.** Phenotypic Prediction of IQ from Each Algorithm in the Training and Test Samples

| Algorithm | r with IQ | Se Lower | Se Upper | P-value |
| --- | --- | --- | --- | --- |
| Lasso Train | 0.380 | 0.349 | 0.410 | <2.2e-16 |
| Ridge Train | 0.476 | 0.447 | 0.503 | <2.2e-16 |
| Shen Train | 0.322 | 0.290 | 0.353 | <2.2e-16 |
| Lasso Test | 0.188 | 0.172 | 0.205 | <2.2e-16 |
| Ridge Test | 0.212 | 0.195 | 0.228 | <2.2e-16 |
| Shen Test | 0.187 | 0.171 | 0.204 | <2.2e-16 |

Note. Phenotypic prediction of IQ by each algorithm, measured as the Pearson's r between brain-based predicted IQ and measured IQ. Training was done in the 3SK sample and testing the 13SK sample. The top three columns represent the algorithm predicting in the sample it was trained in (with LASSO and Ridge using 10-fold cross-validation to guard against overfitting), the bottom three represent the out of sample prediction going from the 3SK to the 13SK sample.

### Predictive features

For the sake of interpretability and to demonstrate the utility of this approach for theory building, we plotted edges included in the Shen et al. model and weights for the ridge and LASSO regression against the Yeo 7 resting state-network parcellation in Figure 3. Due to the large sample size and number of edges, both the Shen et al. procedure and the ridge lack a simple interpretation. In contrast, the LASSO was highly interpretable, as only a handful of edges were needed to account for similar proportions of variance as explained by the Shen et. al. model and the ridge. The LASSO weights were largely negative weights on edges connecting the default-mode network to other networks, i.e. less connectivity between the default-mode network and attention and frontal networks is predictive of higher IQ (by the LASSO output). The edges were anatomically largely default-mode to default-mode edges in and around posterior cingulate cortex.

**Figure 3.**
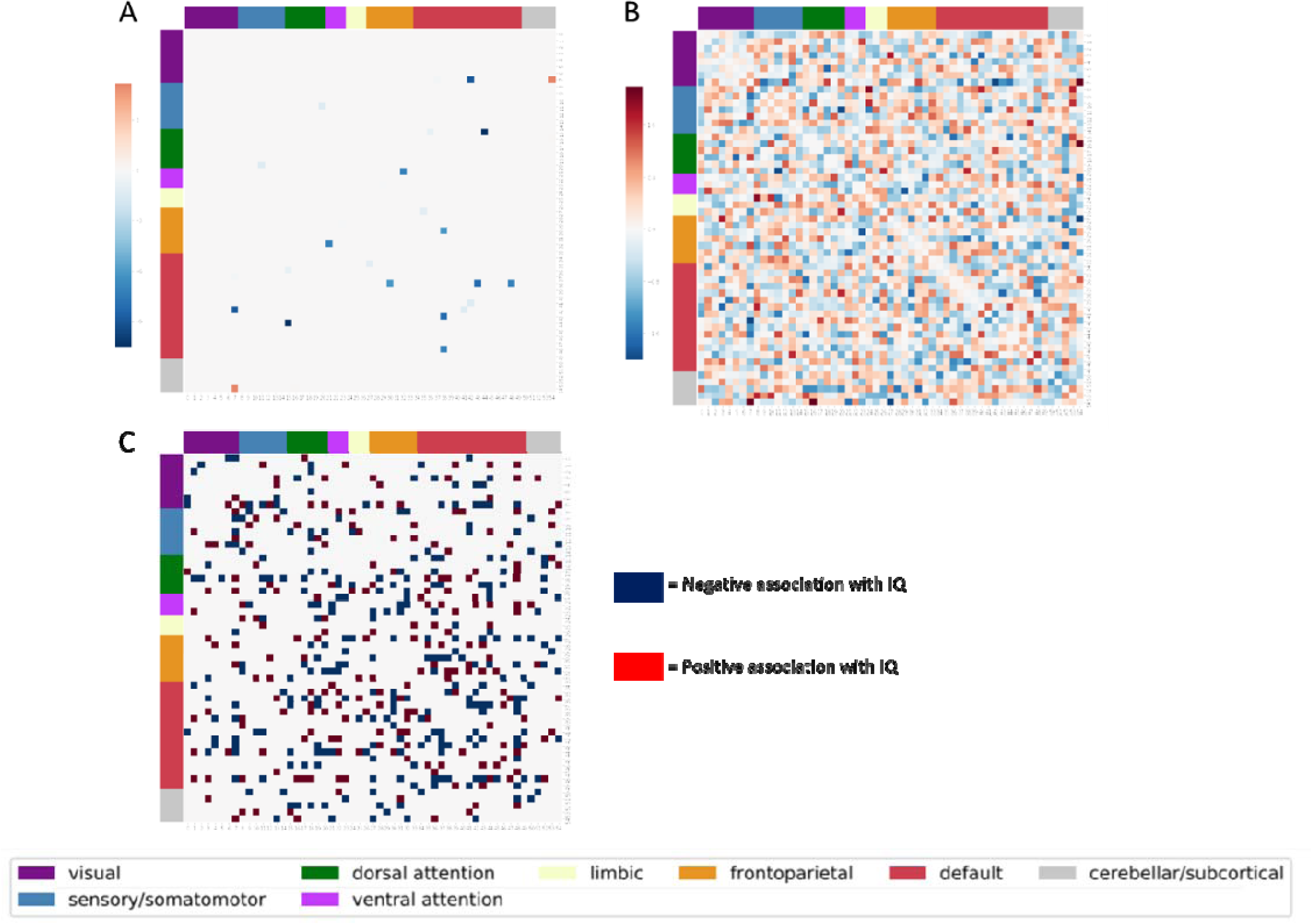
Weights implicated by each predictive modeling technique organized by the Yeo 7 resting state network to increase their interpretability. (A) Weights from the LASSO regression model. (B) Weights by the ridge regression model. (C) Weights that went into the positive (red) and negative (navy blue) sum scores for the Shen et al. procedure.

Because the modeling techniques produced different solutions, we hypothesized that a model using the LASSO selected edges is likely the most predictive, but other edges could be used for prediction beyond the LASSO edges. Comparisons between LASSO and ridge regression models revealed that the specific connections identified by the LASSO model (depicted in Figure 4A) are in fact driving part of the signal to predict IQ, but other edges could be used to effectively (but more weakly) to predict IQ, reconciling the ridge and LASSO results. Further, a repeated *k*-fold cross-validation test revealed that the optimal LASSO and ridge regression models performed better when weights were estimated on LASSO-selected edges only (LASSO: mean difference = .1131, *SD* of difference = 0.0146, *t*_99_ = 4.114, *p* < .0001; ridge: mean difference = .0814, *SD* of difference = 0.0111, *t*_99_ = 4.738, *p* < .0001). Thus, these results indicate that the LASSO-selected edges informed prediction in both models, but that other connected edges are influencing the ridge and Shen et al. procedures and are useful in predicting IQ.

**Figure 4.**
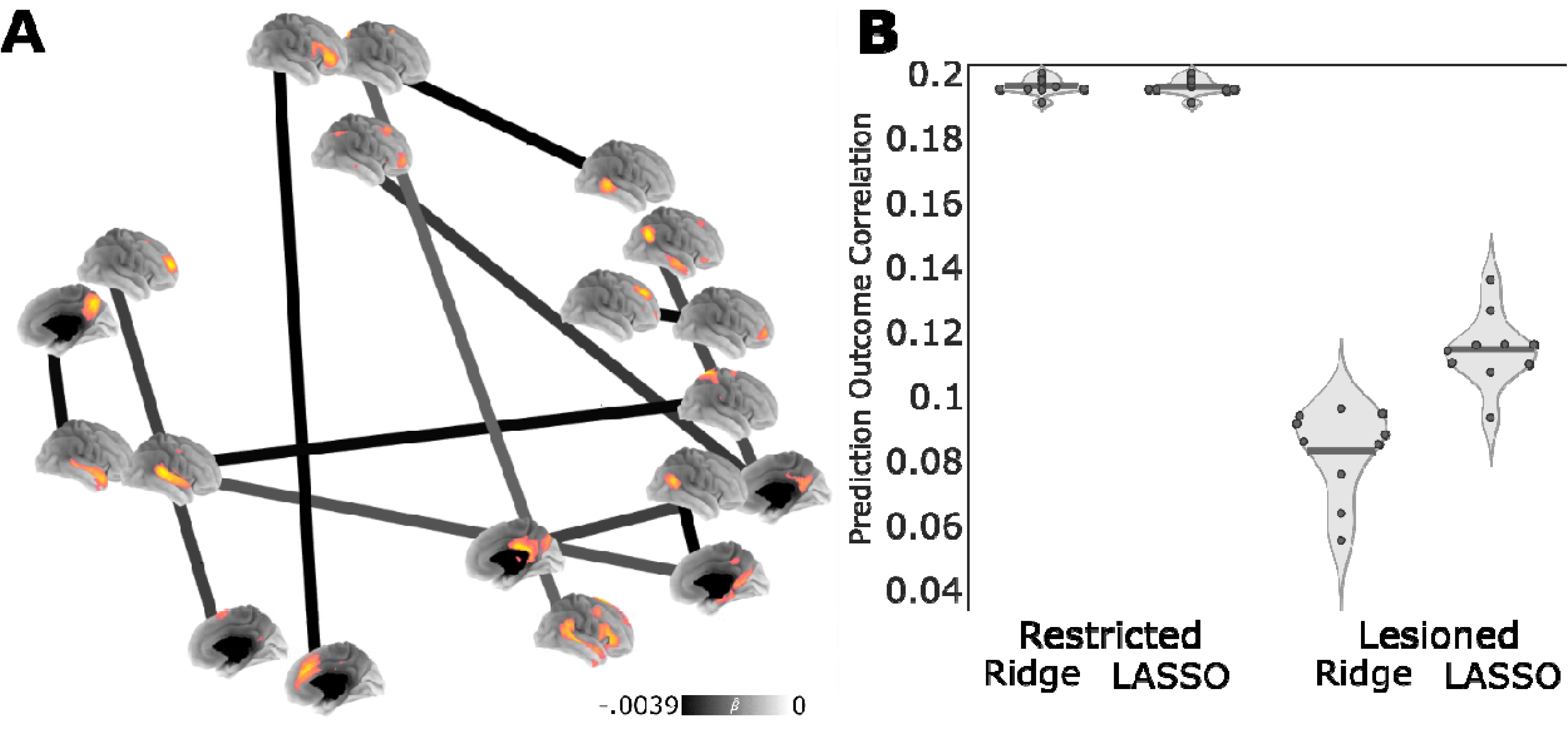
Functional connections necessary for the prediction of IQ. (A) Beta estimates from the LASSO regression are depicted as a graph, where darker edges have more negative weights that indicate greater connectivity is associated with lower IQ. Surface renders depict the spatial distribution of nodes from the UKBiobank connectome, with positive loadings depicted in yellow. (B) Violin plots show the performance of models restricted to the most predictive edges identified by LASSO (i.e., the 12 edges shown in panel A) and models where the same edges were “lesioned” or removed from the full graph. Each point in the plot corresponds to the average correlation between predicted and observed IQ for each of 10 iterations of 10-fold cross-validation. Solid lines indicate the mean across iterations.

### Genetic analyses

#### Univariate heritability

First, to examine the heritability of these scores with these data and check for our multi-system brain models heritability out of sample, we used BOLT-REML to run a univariate heritability analysis of each model’s prediction. All three model predictions were significantly heritable (Table 2), and had similar heritability of approximately 0.15. We also tested the heritability of measured IQ in this sample, which was 0.28 and higher than any predicted model.

**Table 2.** Heritability and Coheritability of Measured IQ and the Prediction of IQ by Each Modeling Procedure

| Edge/Model | Network | Heritability | SE | rG<br>with IQ | SE | rE<br>with IQ | SE | rG CI Corrected |
| --- | --- | --- | --- | --- | --- | --- | --- | --- |
| **Angular Gyrus to PCC** | Default to ventral attention | 0.104 | 0.031 | -0.226 | 0.130 | -0.037 | 0.028 | ±0.361 |
| **Right Frontal Parietal Network Regions and contralateral cerebellum to PCC** | Frontoparietal to ventral attention | 0.070 | 0.030 | -0.353 | 0.167 | -0.008 | 0.027 | ±0.464 |
| **Medial superior occipital to Left middle frontal and temporal structures** | Dorsal attention to default mode | 0.087 | 0.030 | -0.130 | 0.141 | -0.064 | 0.028 | ±0.390 |
| **precuneus, lateral occipital and fusiform to middle temporal Gyrus** | Visual to default mode | 0.102 | 0.030 | -0.252 | 0.133 | 0.004 | 0.028 | ±0.368 |
| **Frontal Pole to Left Frontoparietal network and cerebellum to FP** | Frontoparietal to ventral attention | 0.118 | 0.030 | 0.105 | 0.122 | -0.049 | 0.028 | ±0.339 |
| **Precuneus and PCC to PCC** | Default mode to default mode | 0.038 | 0.029 | -0.310 | 0.241 | 0.009 | 0.027 | ±0.669 |
| **CBPM\*** |  | 0.181 | 0.031 | 0.422 | 0.092 | 0.123 | 0.029 | ±0.256 |
| **Lasso\*** |  | 0.143 | 0.031 | 0.518 | 0.105 | 0.107 | 0.028 | ±0.290 |
| **Ridge\*** |  | 0.155 | 0.031 | 0.576 | 0.100 | 0.116 | 0.028 | ±0.278 |

Note. The heritability of each edge that was significantly associated after Bonferroni correction and the prediction of IQ and coheritability between each edge and each predicted model of IQ with measured IQ. $r_G$ = genetic correlation between the IQ predictor and measured IQ, positive genetic correlation means that the genes that influence stronger connectivity (correlated or anti-correlated) are related to better IQ, and negative means genes that influence connectivity are related to lower IQ. Standard errors (SE) follow their respective statistic. We Bonferroni corrected the confidence interval to the 99.99 % interval to account for multiple testing. PCC= posterior cingulate cortex. $*p < .05$ $r_G$ with brain-based predicted IQ post Bonferroni correction.

#### Genetic correlations between predicted and measured IQ

Next, we estimated the genetic correlation between measured IQ and the prediction of IQ by each method. All models were significantly genetically correlated with measured IQ, with moderate genetic correlations ranging from .422 - .576 (Table 2). The genetic correlations followed the same pattern as the phenotypic associations: The ridge was most genetically correlated with measured IQ. Importantly, this means that *more than half of the genetic variance influencing IQ* can be captured by ridge regression and LASSO regression brain-based predicted IQ scores.

Considering that various measured IQ cohorts from Savage et al.^26^ meta-analysis found an average rG of .67, our brain-based predicted IQ variable is approaching similar utility to one of these GWAS meta-analysis cohorts.

Next, we compared genetic correlations of measured IQ with the six single edges associated with IQ in Figure 3A to genetic correlations from whole brain models. Table 2 shows the results and the 6 edges significantly phenotypically associated with IQ, post Bonferroni correction. All single edges were theoretically relevant. The top associated edge was a connection involving the angular gyrus, which smaller samples have shown to be associated with IQ in the literature^33^. Several edges contained the posterior cingulate cortex, part of the default network which is known to decrease in activation during demanding, externally directed cognitive tasks^34^, and thus individuals with greater connectivity in the posterior cingulate may be showing less engaged task-positive network activity and lower IQ. Notably, the three brain models were nominally more heritable than the single brain edges. After correction for multiple tests, no individual edge was significantly genetically associated with measured IQ, in line with past studies reporting null effects for brain-behavior genetic correlations (9 tests). Nominally, all genetic correlations between single edges and measured IQ were lower than genetic correlations between predicted and measured IQ. This is in line with our hypothesis that multi-system predictive approaches will be more useful for linking genes, brain and behavior as they are more heritable and show higher genetic correlations than single measures of the brain.

#### GWAS of brain-based predicted IQ

To test if brain-based predicted IQ is a useful GWAS proxy, we ran a GWAS with the prediction from each machine learning algorithm as the phenotype and compared it to a GWAS of measured IQ in the same individuals. No GWAS found significant SNPs above genome-wide p-value significance (see Figure 5 for Manhattan plots of each of the 4 traits). This suggests very large GWAS samples are going to be needed to detect significant variants of predictive models, which is line with sample size calculations needed for GWAS studies^35^. We used the QQ Plots of estimated vs. expected p-values to probe the power from each of these models’ predictions and measured IQ (Figure 6). Both the LASSO and the ridge regression models had similar polygenic signals as measured IQ. However, the Shen et al. procedure led to the greatest deviation from expected p-values. Arguably, the Shen method may offer more power to detect variants than a standard GWAS phenotypes, but this is an empirical question and here did not lead to increased GWAS discovery.

**Figure 5.**
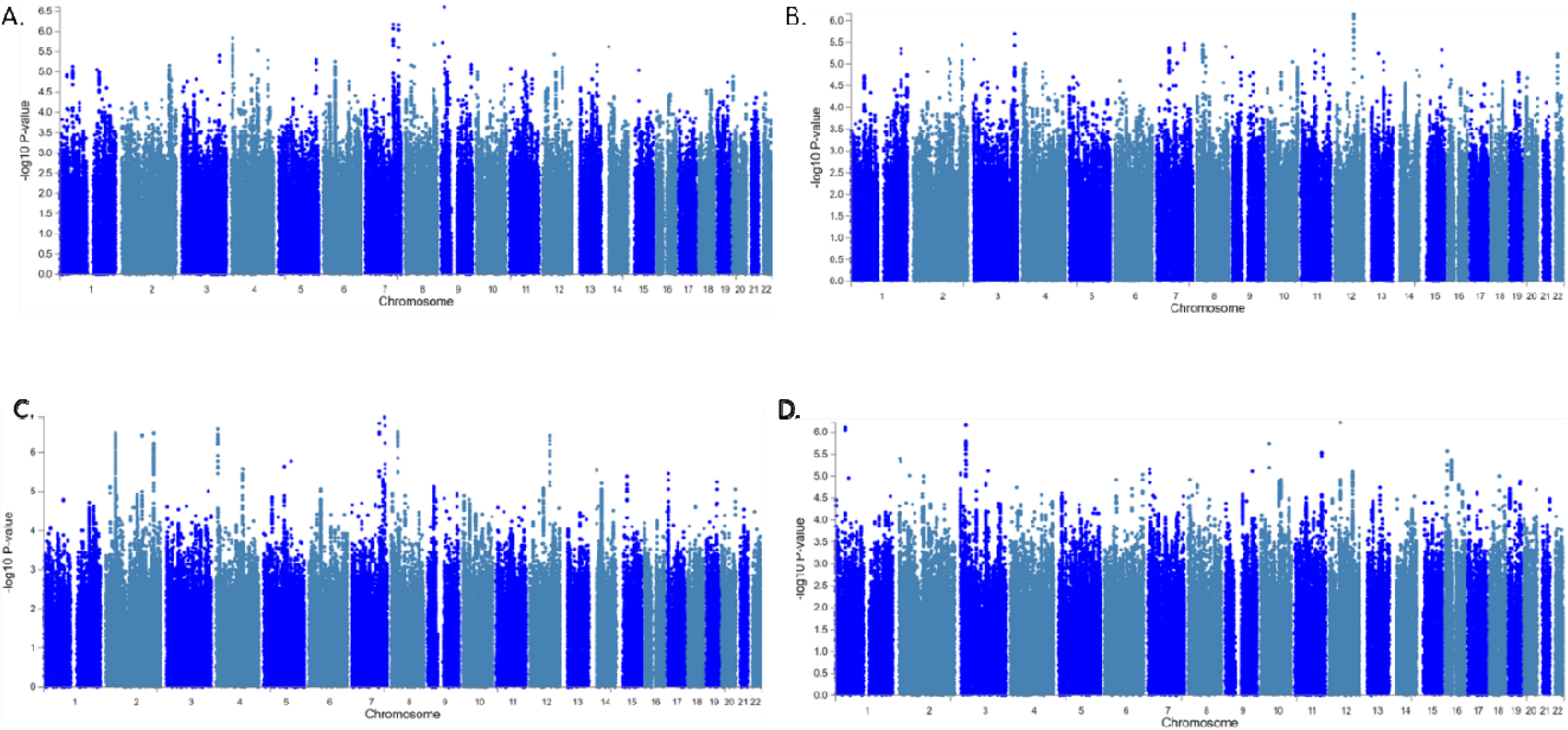
Manhattan plots for genome-wide association discovery across the three predictive models (A is LASSO, B is ridge, and C is the Shen et al. procedure) and measured IQ (D) in the same individuals from the 13SK sample. P-values of each SNP are organized by chromosome on the x-axis and by the -log10 of the p-value on the y axis. No p-values were significant below genome-wide discovery correction (p < 5e-8).

**Figure 6.**
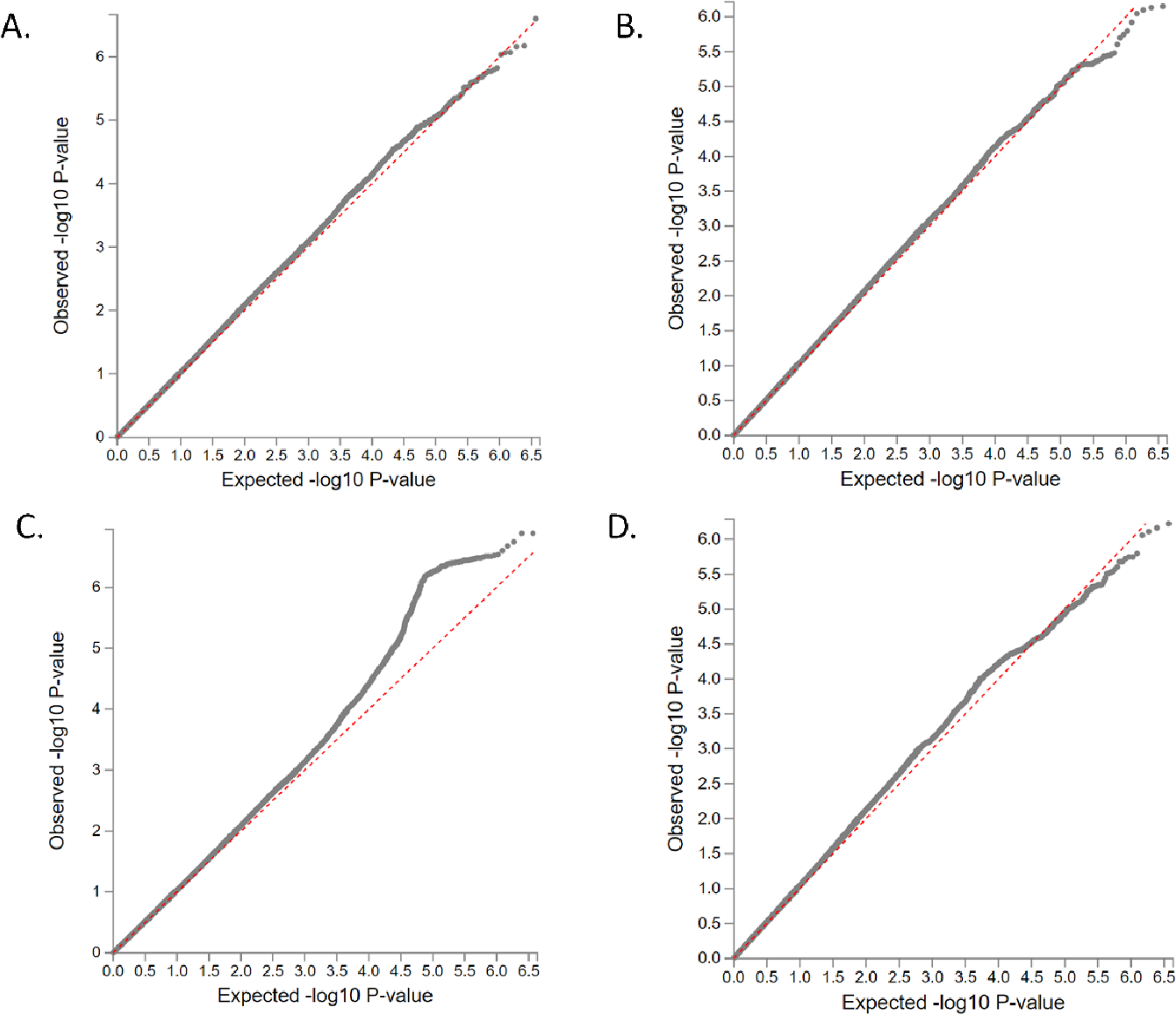
QQ-plots for the p-values from the genome-wide association discovery across the 3 predictive models (A is LASSO, B is ridge, and C is the Shen et al. procedure) and measured IQ (D) in the same individuals from the 13SK sample. P-values are the dotted line plotted based on their expected -log 10 p-value on the x-axis and by their observed – log 10 p-value on the right access. The dashed line is the expected p-value distribution under the null model. Deviation above the line represents signal more significant associations than expected.

#### Genetic correlations between brain-based predicted IQ and correlates of IQ

To see if brain-based predicted IQ endophenotypes also were genetically correlated with phenotypes genetically correlated with measured IQ, we conducted LD score regression genetic correlations of each model predictions with 8 key traits of interest: years of education, college completion, Childhood IQ, depressive symptoms, autism spectrum disorder, height, infant head circumference, and age at first birth. We compared these genetic correlations between brain-based predicted IQ and traits of interest to genetic correlations of measured IQ (in the same individuals as the predictive IQ scores were drawn from) and these same 8 key traits of interest. All three predictive models of IQ were highly genetically correlated with one another, so the genetic correlations do not change appreciably based on the method of deriving the model (Ridge-Lasso rG = .946, CI = ± .033; Ridge-Shen rG = .902, CI = ± .031; Lasso-Shen, rG = .843, CI = ± .044). Table 3 shows the results. IQ proxies were significantly genetically correlated with all IQ correlates, except Autism Spectrum Disorder. Interestingly, the brain-based predicted models were genetically associated with early life developmental outcomes. Notably, genetic predisposition to high IQ-predictive brain model responses was associated with genetic predisposition to higher educational attainment (rG = 0.353 and 0.420 for years schooling/college), large infant head circumference (rg = 0.600) and childhood IQ (rG = .453). Thus, the brain models captured genetically linked variation shared with other IQ-related early life traits. Essentially, the genes that confer a genetic advantage in early life also confer stronger IQ-related brain responses, giving some credence the stability of the genes captured by fMRI connectivity in mid-life.

**Table 3.** Genetic Correlations Between Brain-based Predicted IQ and Known Genetic Correlates of IQ

| **IQ Model with**<br>IQ-correlated trait | **rG** | **se** | **Z** | **p-value** |
| --- | --- | --- | --- | --- |
| **Ridge with** |  |  |  |  |
| Years of schooling | 0.353 | 0.082 | 4.311 | <0.001 |
| College completion | 0.420 | 0.120 | 3.493 | 0.001 |
| Depressive symptoms | -0.456 | 0.139 | -3.285 | 0.001 |
| Age of first birth | 0.289 | 0.090 | 3.206 | 0.001 |
| Childhood IQ | 0.453 | 0.154 | 2.941 | 0.003 |
| Infant head circumference | 0.638 | 0.222 | 2.878 | 0.004 |
| Height | 0.023 | 0.062 | 0.370 | 0.711 |
| Autism spectrum disorder | 0.246 | 0.131 | 1.877 | 0.061 |
| **Lasso with** |  |  |  |  |
| Years of schooling 2016 | 0.381 | 0.113 | 3.359 | 0.001 |
| College completion | 0.397 | 0.148 | 2.687 | 0.007 |
| Depressive symptoms | -0.440 | 0.176 | -2.495 | 0.013 |
| Age of first birth | 0.238 | 0.111 | 2.146 | 0.032 |
| Childhood IQ | 0.573 | 0.232 | 2.470 | 0.014 |
| Infant head circumference | 0.747 | 0.286 | 2.614 | 0.009 |
| Height | 0.011 | 0.074 | 0.144 | 0.886 |
| Autism spectrum disorder | 0.263 | 0.170 | 1.550 | 0.121 |
| **Shen with** |  |  |  |  |
| Years of schooling | 0.344 | 0.079 | 4.356 | <0.001 |
| College completion | 0.391 | 0.121 | 3.230 | 0.001 |
| Depressive symptoms | -0.428 | 0.148 | -2.891 | 0.004 |
| Age of first birth | 0.248 | 0.095 | 2.613 | 0.009 |
| Childhood IQ | 0.298 | 0.159 | 1.871 | 0.061 |
| Infant head circumference | 0.676 | 0.242 | 2.793 | 0.005 |
| Height | 0.086 | 0.072 | 1.192 | 0.233 |
| Autism spectrum disorder | 0.195 | 0.127 | 1.529 | 0.126 |
| **Measured IQ with** |  |  |  |  |
| Years of schooling 2016 | 0.618 | 0.062 | 9.936 | <0.001 |
| College completion | 0.653 | 0.087 | 7.481 | <0.001 |
| Depressive symptoms | -0.239 | 0.087 | -2.740 | 0.006 |
| Age of first birth | 0.446 | 0.072 | 6.172 | <0.001 |
| Childhood IQ | 0.780 | 0.151 | 5.184 | <0.001 |
| Infant head circumference | 0.438 | 0.151 | 2.895 | 0.004 |
| Height_2010 | 0.182 | 0.051 | 3.567 | <0.001 |
| Autism spectrum disorder | 0.136 | 0.095 | 1.436 | 0.151 |

Note. Genetic correlations (rG) between brain-based predicted IQ and genetic correlates of IQ, compared to genetic correlations with measured IQ. Genetic correlations were estimated using
LD Score regression from summary statistics were obtained in GWAS. Data on IQ genetic correlates are from summary statistics of major GWAS consortia.

IQ was more strongly and significantly genetically correlated with educational attainment and (nominally) with, height and age at first birth, to be expected from a direct (vs. proxy) measure. However depressive symptoms, and infant head circumference were more correlated with brain-based predicted IQ than measured IQ (nominally).

### Utility of Predictive Models Across Samples

Finally, we conducted post-hoc analysis to see what sample sizes would be needed to use in development of CBPM in the UKBiobank parcellation. Here, we also allowed the p-value threshold of the Shen et al. procedure to vary. Figure 7 shows the results of learning curves training the S13K and predicting in the S3K sample. More than 1000 individuals are needed to reduce overfitting of the machine learning procedures (at least within this sample and parcellation). In line with results from genetic predictive models^36^, the Shen et al. count model seems to perform the best in smaller sample sizes (less overfitting) but becomes less useful as the sample sizes increase. The LASSO regression outperformed the ridge regression with small samples and performed worse than the ridge but with similar prediction with larger sample sizes. Thus, in this study LASSO regression offered interpretability and was a useful predictor across a range of sample sizes, meaning that LASSO is a well-balanced procedure for future work. Finally, as sample sizes became large, the ridge prediction slightly outperformed the LASSO.

**Figure 7.**
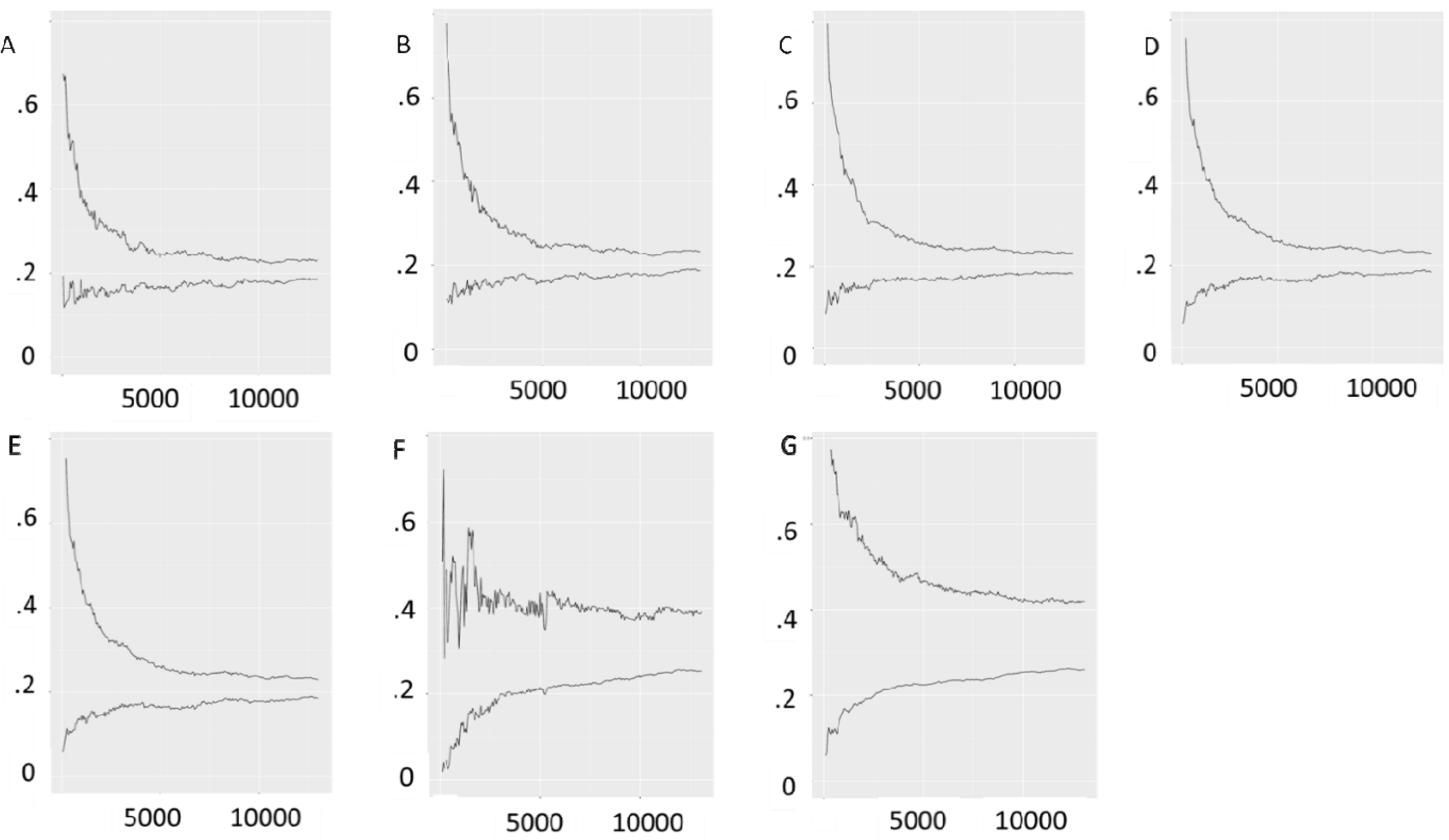
Learning curves for increasing sample sizes of predictive models of intelligence. The top line is the curve in the training set as sample sizes increases in the training set. The bottom line is prediction in test set of 3000 individuals as training sample size increasing. (A) Shen et al. p-value threshold =.005. (B) Shen et al. p-value threshold =.01. (C) Shen et al. p-value threshold =.05, value that is often chosen. (D) Shen et al. p-value threshold =.1. (E) Shen et al. p-value threshold =1. (F) LASSO regression. (G) Ridge regression.

## Discussion

We found that predictive modeling was an effective way to generate endophenotypes. We demonstrated the application of multi-system brain predictive models as GWAS proxies for discovery and theory building. We also presented novel innovations for estimation of the scores phenotypically with CBPM procedures and compared different approaches for developing these models using UKBiobank resting state connectivity data. LASSO regression developed endophenotypes that were interpretable and predicted more than half of the genetic variance underlying IQ in out-of-sample predictions. The Shen et al. procedure seemed to offer the most power to detect polygenic signal, based on deviation from expected p-values. Conclusions for this approach are presented below.

### Multi-system brain endophenotypes and unmeasured variance

We established that multi-system brain predictive models are useful endophenotypes and should be applied to work in the future. One application of this approach could be to expand the range of possible behavioral phenotypes to conduct GWAS of traits in these larger samples. For example, in the imaging literature there are demonstrations where brain models of dichotomous traits with small samples are combined with brain models from continuous measured mechanisms to increase power, which has led to increased predictive accuracy for fibromyalgia (dichotomous) using a pain sensitivity model (continuous)^37^. This could expand the number of GWAS phenotypes and direct efforts to even predict particular mechanisms underlying these traits.

When endophenotypes were first purposed as useful biomarkers, it was believed they would play a key role in gene discovery^5^. While they were largely unsuccessful at this goal, they have transitioned into useful measures for understanding underlying psychological variability^7^.

As many CBPM models reflect behavioral phenotypes close to brain and mechanisms in behavior (i.e. sustained attention in ADHD), GWAS of these models may offer a path forward in understanding the interplay of behavioral and genetic mechanisms in downstream health and wellness outcomes.

While it is often argued that predictive modeling approaches are difficult to interpret, plotting the weights can lead to insights into the brain systems underlying cognitive and behavioral phenotypes. In line with modern endophenotype approaches, we also demonstrate how prediction across the connectome is useful for theory building; and align our approach with the common Yeo 7 parcellation to establish functional implications of these associations. Specifically, using LASSO regression, we found that default-mode to other network connections had high utility in predicting IQ, in line with the current literature^33, 34^, and also implicated 6 edges that were associated via the standard approach of univariate tests with multiple correction. We also went beyond these simple associations by discovering signal for 12 edges (with LASSO) and demonstrating that connectivity patterns beyond these 12 are associated with IQ. Thus, multi-system approaches give complementary evidence to univariate approaches, and extend to more complex patterns.

Further, by comparing methods for predictions, we were able to see how the signal for predicting IQ from the connectome was (1) driven by a handful of interpretable edges and (2) related to larger multi-system brain patterns. These results are in line with imaging literature that argues IQ relies on modularity, or dynamic reprocessing and repurposing existing network structures, rather than being driven by the strength of key nodes in the network^38^. Thus, multi-system brain predictive approaches align with modern endophenotype research and brain discovery. Importantly, our approach does not limit discovery to single brain regions; more edges were weighted in the LASSO than were associated via correction after the univariate tests of association, and the full weighted model showed much stronger genetic associations than any single edge, giving us a better picture of the multivariate pattern underlying IQ.

Further, CBPMs can be used to find genetic correlations with traits genetically related to the trait of interest. In this study, traits genetically correlated with IQ were genetically correlated with the brain-based predictions of IQ. Interestingly, infant head circumference and depressive symptoms were more genetically correlated with brain-based predicted IQ than measured IQ. This may mean that the association between IQ and connectivity is strongly mediated by these phenotypes or vice versa. For example, it appears that the main edges driving the prediction of IQ (according to the LASSO output) were in the default mode network, which is thought to play a key role in depressive symptoms^39^. Therefore, it is likely that default mode connectivity genetics is influencing the overlap between intelligence and depressive symptoms.

### Predictive modeling insights from this study

Based on patterns of results, we recommend that CBPMs utilize large samples. When comparing the different methods, the standard CBPM (Shen et al. 2017) seemed to find solutions that worked effectively in smaller samples, with adjustments on the p-values only providing marginal change to the model performance. LASSO regression was useful across a range of sample sizes after samples exceeded ∼1,000 individuals. Ridge regression seems to require very large sample sizes to avoid overfitting. The Shen sum score approach outperformed other approaches in smaller samples, but regularization in larger samples mirrored results from the statistical genetics literature on polygenic risk scores^40^. A key difference is that much smaller sample sizes are needed for the same level of prediction in multi-system brain models than polygenic risk scores (only a couple thousand compared to 10’s to 100’s of thousands).

There are many neuroimaging modalities across the functional and anatomical literature and many possible levels of analysis via different parcellations that can be used for multi-system brain predictive modeling. This study establishes the utility of multi-system brain modeling with the UKB functional connectivity parcellation and CBPM procedures. The procedures here predicted some, but not all, of the genetic variability underlying IQ. Though it would be outside the scope of any one manuscript to test all possible ways of predicting an outcome and its genetic associations, in the future, many other sources of data and brain parcellations may be used to expand on this method. Woo et al.^41^ offer a procedure for selecting the best model that involves first testing different models in the discovery sample, selecting the one that is predictive (and makes theoretical sense) and then holding out a validation dataset just for that model in particular. Using procedures like this that have been established in imaging may improve use of endophenotypes in the future.

### Limitations

There were a number of limitations to the study. First, our training and validation sets were closely aligned as they are both part of the UKBiobank. It is possible that these results may generalize poorly to other parcellations or subgroups (like younger cohorts) that were not used in generation of the UKBiobank parcellation. Consideration of brain parcellation and feature engineering is an important step in choosing predictive models, and this work does not argue against the importance of those considerations.

In our gene finding we focused on common variants (MAF > .01) and highly probable variants (INFO > . 95). We did so because we were running several GWAS and we wanted to conduct reasonable genome-wide screening, but also reduce the computational burden. This choice means, however, that we cannot speculate about how rare variants are likely influencing multi-system brain models.

### Conclusions

Multi-system brain models will be useful endophenotypes for neuropsychological outcomes in the future. Connectivity-based models performed well for predicting IQ in this study, predicting about half the genetic variance underlying IQ. Future expansions should consider other neuroimaging modalities as ways to further improve these scores.

## Acknowledgments

The Authors would like to acknowledge NIH grants MH063207, DA046064, DA042742, T32MH016880-38, and T32DA007261-28. We would also like Cold Spring Harbor Laboratory for hosting the preprint of this manuscript on Bioarchive. This research has been conducted using the UK Biobank Resource under Application Number 24795.

